# The Ensembl COVID-19 resource: Ongoing integration of public SARS-CoV-2 data

**DOI:** 10.1101/2020.12.18.422865

**Authors:** Nishadi H. De Silva, Jyothish Bhai, Marc Chakiachvili, Bruno Contreras-Moreira, Carla Cummins, Adam Frankish, Astrid Gall, Thiago Genez, Kevin L. Howe, Sarah E. Hunt, Fergal J. Martin, Benjamin Moore, Denye Ogeh, Anne Parker, Andrew Parton, Magali Ruffier, Manoj Pandian Sakthivel, Dan Sheppard, John Tate, Anja Thormann, David Thybert, Stephen J. Trevanion, Andrea Winterbottom, Daniel R. Zerbino, Robert D. Finn, Paul Flicek, Andrew D. Yates

## Abstract

The COVID-19 pandemic has seen unprecedented use of SARS-CoV-2 genome sequencing for epidemiological tracking and identification of emerging variants. Understanding the potential impact of these variants on the infectivity of the virus and the efficacy of emerging therapeutics and vaccines has become a cornerstone of the fight against the disease. To support the maximal use of genomic information for SARS-CoV-2 research, we launched the Ensembl COVID-19 browser, incorporating a new Ensembl gene set, multiple variant sets (including novel variation calls), and annotation from several relevant resources integrated into the reference SARS-CoV-2 assembly. This work included key adaptations of existing Ensembl genome annotation methods to model ribosomal slippage, stringent filters to elucidate the highest confidence variants and utilisation of our comparative genomics pipelines on viruses for the first time. Since May 2020, the content has been regularly updated and tools such as the Ensembl Variant Effect Predictor have been integrated. The Ensembl COVID-19 browser is freely available at https://covid-19.ensembl.org.

## INTRODUCTION

Over the past twenty years, multiple zoonotic respiratory diseases caused by coronaviruses have been identified. Examples include the SARS epidemic caused by severe acute respiratory syndrome coronavirus (SARS-CoV) in 2003 and the Middle East respiratory syndrome coronavirus (MERS-CoV) outbreak in 2012. Both belong to the *betacoronavirus* genus and are believed to have originated in bats with an intermediary animal host before transmission to humans^1^.

<u>T</u>he SARS-CoV-2 virus responsible for the current COVID-19 pandemic is also a *betacoronavirus*, with a 29,903-nucleotide positive-strand RNA genome encoding ~30 known and hypothetical mature proteins. The first open reading frame (ORF), representing approximately 67% of the entire genome, encodes 16 non-structural proteins (nsps). The remaining ORFs encode accessory proteins and four major structural proteins: spike surface glycoprotein (S), small envelope protein (E), matrix protein (M) and nucleocapsid protein (N).

Genomic sequencing has played a crucial role in understanding the mechanisms, spread and evolution of this virus. In the UK alone, at the time of writing, close to 5% of all reported infections each week were being sequenced (COG-UK, January 2021: https://www.cogconsortium.uk/wp-content/uploads/2021/02/COG-UK-geo-coverage_2021-02-01_summary.pdf) and this trend is likely to grow. Established genomic resources, such as Ensembl, have been able to leverage these data and bring them to new and existing user communities supporting research leveraging the rapidly emerging SARS-CoV-2 data landscape.

Ensembl^2^,^3^ was launched to capture data from the Human Genome Project and has since developed into a large scale system for generating, integrating and disseminating genomic information. The COVID-19 pandemic presented new challenges related to presenting SARS-CoV-2 annotation and data within Ensembl. Meeting these, we launched the Ensembl COVID-19 browser (https://covid-19.ensembl.org) in May 2020 using concepts and workflows that enable rapid update cycles to react quickly in the face of new data and potential future outbreaks.

## NEW ENSEMBL COVID RESOURCE

### Reference assembly and a new gene annotation

The SARS-CoV-2 sequence represented in Ensembl (INSDC accession GCA_009858895.3, MN908947.3) is the viral RNA genome isolated from one of the first cases in Wuhan, China^4^. It is widely used as the standard reference and has been incorporated into other resources such as the UCSC SARS-CoV-2 genome browser^5^. This assembly was imported from the European Nucleotide Archive (ENA) into an Ensembl database schema with minor modifications to software regularly used to integrate assemblies from the ENA into Ensembl.

To enable the correct annotation of SARS-CoV-2, the Ensembl gene annotation methods^6^ were adapted to reflect the biology of the virus. To identify protein coding genes, we aligned SARS-CoV-2 proteins from RefSeq^7^ to the genome using Exonerate^8^. A challenge for annotation is that the first and largest ORF can result in either non-structural proteins nsp1-11 (ORF1a) or in nsp1-nsp10 and nsp12-nsp16 (ORF1ab) via an internal programmed translational frameshift^9^. Exonerate handles this ribosomal slippage by inserting a gap in the alignment and thus allowing the annotation of the full ORF1ab locus. Our modified annotation methodology then removes the artificial gap to represent the slippage frameshift as an RNA edit and ensures a biologically accurate representation of the locus and product.

Our annotation approach was tested on 90 additional SARS-CoV-2 assemblies retrieved from the ENA. We assessed alignment coverage and percentage identity of the resultant gene translations to verify accuracy and consistency. In all cases, full length alignments were observed and average amino acid percentage identity across all genes in most assemblies were 99.9% or 100% (one assembly had 99.81% identity). These results demonstrate that our annotation approach is able to scale consistently to larger volumes of viral data.

In addition to generating a fully integrated Ensembl gene annotation, we also imported the gene set submitted to the ENA with the reference sequence by the Shanghai Public Health Clinical Centre. As shown in figure 1, both the submitted (blue) and the Ensembl gene annotations (red) can be viewed simultaneously on the browser. The submitted gene annotation is displayed as a separate annotation track, accessed under the ‘Genes and transcripts’ heading after clicking on ‘Configure this page’ in the left-hand menu. The major difference between the annotations is that the submitted annotation does not include the short form ORF1a or ORF7b.

**Figure 1:**
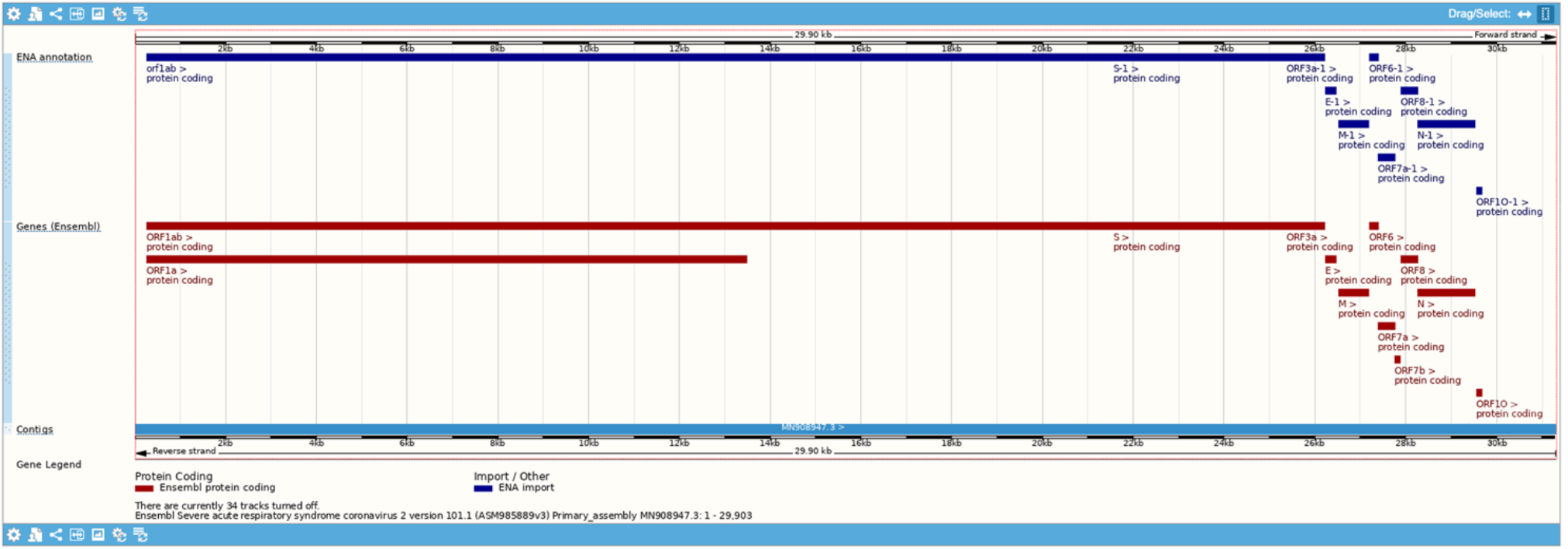
A comparison of the Ensembl gene set and the gene set submitted to the ENA by the Shanghai Public Health Clinical Centre for the SARS-CoV-2 reference assembly

### Comparison of SARS-CoV-2 with 60 other *Orthocoronavirinae* genomes

We used Cactus^11^ to align SARS-CoV-2 to 60 publicly available virus genomes from the *Orthocoronavirinae* subfamily. The results showed 78% of the SARS-CoV-2 genome aligned with at least one other genome and 35% of the genome aligned with the complete set of *Orthocoronavirinae* genomes. The multiple sequence alignment gives evolutionary context for each region of the genome and is a powerful method to explore functionality. For example, comparative genomics information such as this can be used for analyses such as a recent comparison of the gene sets <u>of</u> 44 complete *Sarbecovirus* genomes suggest<u>ing</u> both a potentially novel alternate frame gene ORF3c and that ORF10, ORF9c, ORF3b and ORF3d are unlikely to be protein coding^10^.

The alignment coverage (see figure 2) represents the number of genomes aligned to a given reference genomic position and is distributed heterogeneously across the SARS-CoV-2 genome. An immediate observation is that the central region of the genome (starting from ~7.1kb and ending at 21.3kb), including a significant segment of the 3’ part of ORF1a, is highly shared across the *Orthocoronavirinae* subfamily. This indicates that the non-structural proteins encoded by this region (nsp3 - nsp16) likely originate from the *Orthocoronavirinae* ancestral genome. Conversely, both ends of the SARS-CoV-2 genome have very low alignment coverage and are only shared with closely related viruses. As a further demonstration of the utility of the alignment coverage, we focused in on the genomic region encoding for the SARS-CoV-2 spike protein (figure 2). The spike protein has two subunits: S1 which binds to the host cell receptor angiotensin-converting enzyme 2 (ACE2) and S2, which is involved in membrane fusion. The region of the S ORF encoding for the S2 subunit of the spike protein clearly displays a high alignment coverage while the region encoding for the S1 subunit has large portions that are shared only by one other related genome. This demonstrates the dramatic difference in conservations between the S1 and S2 subunits.

**Figure 2:**
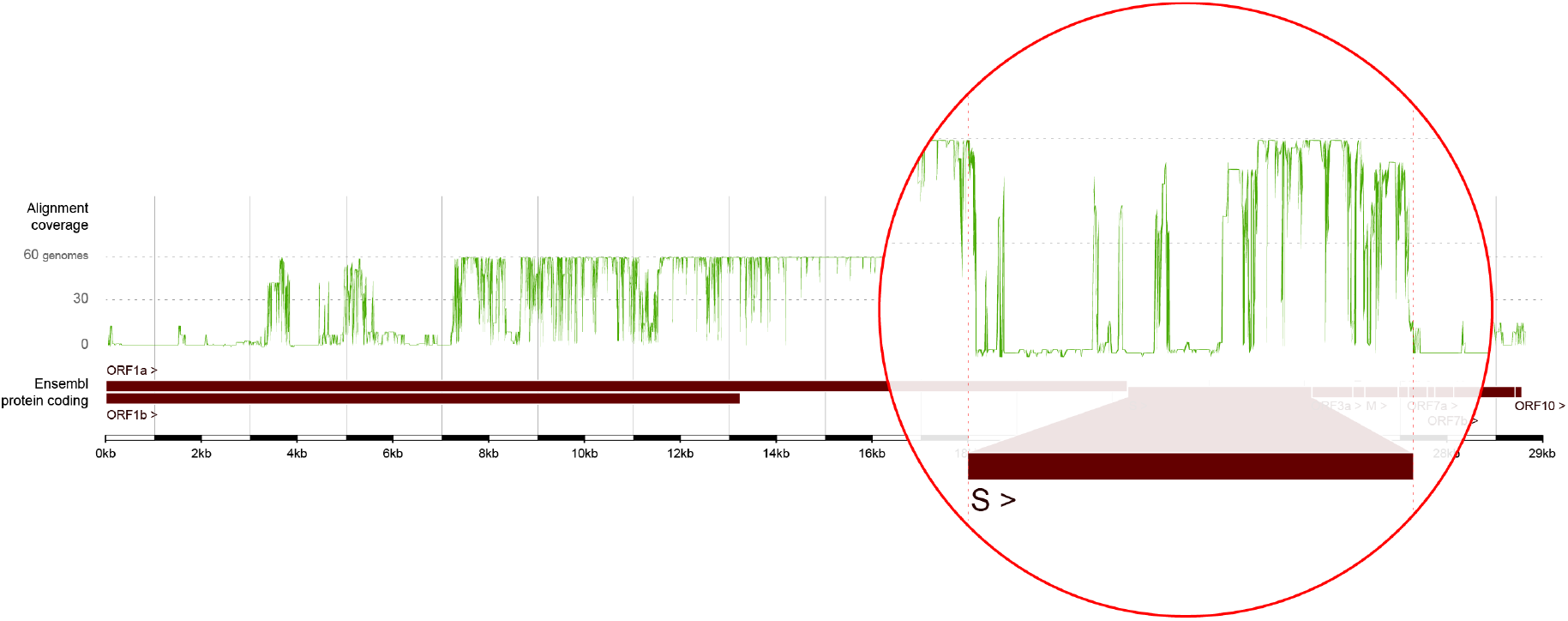
Alignment coverage across the SARS-CoV-2 reference genome based on a multiple sequence alignment with 60 other *Orthocoronavirinae* genomes. The green plot of alignment coverage shows that the central region of the genome is highly shared across the subfamily, while the ends are generally shared only with closely related viruses. The region encoding for the spike protein S has been highlighted within the red circle showing the difference between the low alignment coverage of the upstream S1 subunit and the high coverage of the downstream S2 subunit.

Additionally, we applied our gene tree method^12^ to group the protein coding genes into families and to predict orthologous and paralogous relationships between genes. These results will be incorporated into the COVID-19 resource in Q2 2021.

### Genetic variation data

Analysis of genetic variants of viral genomes is important for understanding the spread of infection across different geographic regions. We display 6,134 sequence variants for SARS-CoV-2 and show their regional frequency distributions alongside predicted molecular consequences calculated by the Ensembl Variant Effect Predictor (VEP)^13^. The variants on our site are derived from overlapping sample sets produced by two groups and a small collection of variants of special interest.

One set comes from the Nextstrain project which creates phylogenetic trees for tracking pathogen evolution based on virus subsamples^14^. We converted their SARS-CoV-2 data release from 08-04-2020 to VCF for integration into our system and display frequency distributions by country and Nextstrain-inferred clade.

The second variant set comes from the ENA team, who developed a LoFreq-based^15^ pipeline to call variants from SARS-CoV-2 sequence data sets submitted to their archives. LoFreq reports the proportion of each variant seen in a sample from an individual. For simplicity, we represent only the alleles seen in each sample and not the proportions estimated. Variants were called for each host sample individually and, to provide a more accurate estimation of the frequency of each allele across the entire sample set, it is assumed that sites at which a variant was not called in a sample match the reference genome used in the Ensembl COVID-19 browser. We currently display ENA’s variant data from 17-08-2020 and have applied strict filters to reduce the proportion of lower confidence sites. Specifically, we have not included variants from sequence data sets with more than 40 calls and we have removed variants where no sample has a frequency of 20% or more for the non-reference allele and variants where all samples show strand bias.

Some sites are annotated as a further guide to quality. For example, variants seen in more than one sample in either set have an evidence status of ‘Multiple observations’ and variants at sites recommended for masking by De Maio *et al*^16^ have a flag of ‘Suspect reference location’. Variants can be displayed as three separate tracks in the genome browser: those from ENA, those from Nextstrain and those observed in more than one sample in either project as shown in figure 3.

**Figure 3:**
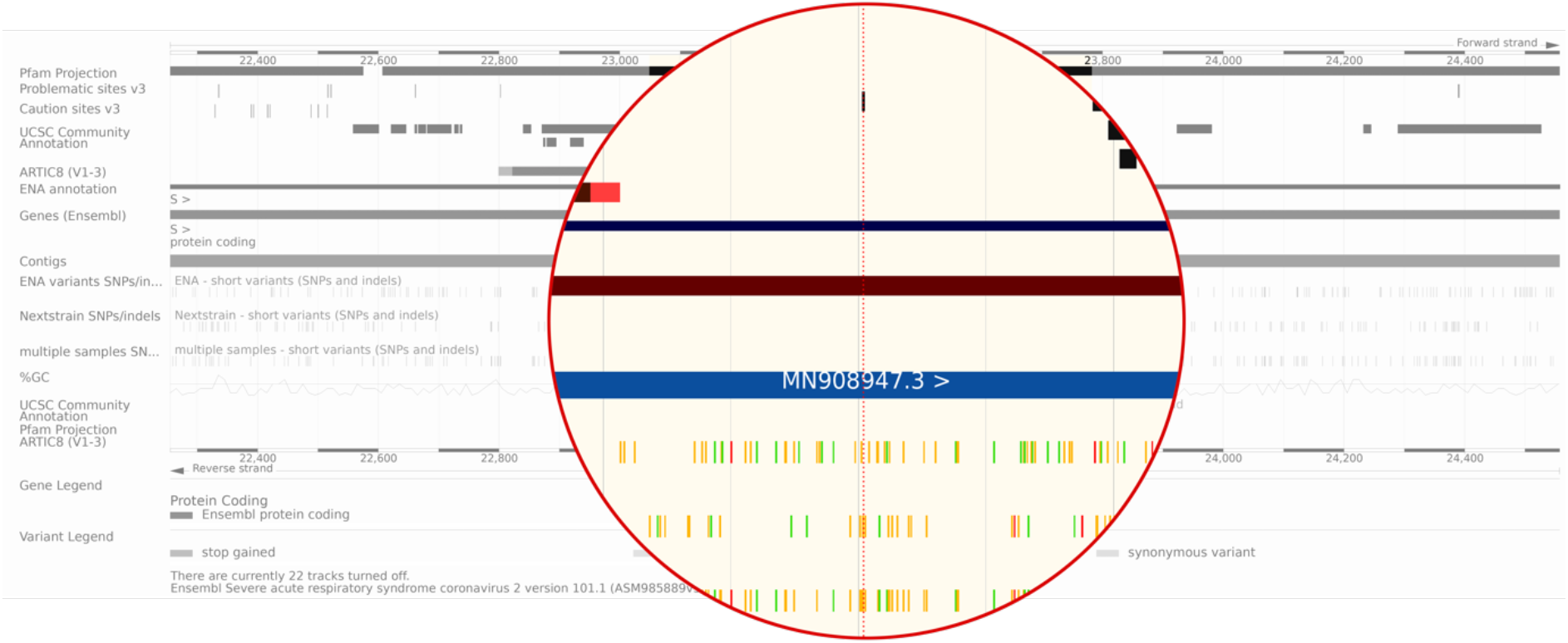
The browser with several tracks turned on and highlighting a substitution flagged up early on in the UCSC community annotation at position 23403 (D614G) in the S spike glycoprotein gene. Due to the prompt nature of community driven annotation, this data was available on our browser as soon as the annotation appeared in a preprint. It is labelled as a common missense mutation in SARS-CoV-2 with a notably high difference in resulting isoelectric point (D->G). Pachetti *et al* (2020) looked at 220 genomic sequences obtained from the GISAID database and characterised 8 novel recurrent mutations; the one at 23403 is one of them. Many studies now show that this particular missense mutation in the spike protein is predominantly observed in Europe^23^; patterns that can also be seen in the variation data we host.

We also display a set of variants which were reported as a tracking priority by the COVID-19 Genomics UK Consortium (COG-UK, https://www.cogconsortium.uk/) in December 2020. This includes 17 variants from the rapidly spreading B.1.1.7 strain (https://virological.org/t/preliminary-genomic-characterisation-of-an-emergent-sars-cov-2-lineage-in-the-uk-defined-by-a-novel-set-of-spike-mutations/563) and four variants from the mink associated strain. The D614G, A222V and N439K mutations associated with an effect on transmissibility, a fast growing lineage and increased binding affinity to the ACE2 receptor^17–19^, respectively, have also been included. We extracted the gene and protein change information from the reports and used the Ensembl VEP to map these descriptions to genomic coordinates and create a VCF file, which was then loaded into the Ensembl database with associated phenotype information. These variants can be viewed as a ‘COG-UK priority mutations’ track alongside the gene annotations.

### Integration of data from other resources

To enrich the SARS-CoV-2 genome annotation we aligned and integrated data from several external repositories in a similar manner to other genomes available in Ensembl.

Specifically, we aligned Rfam^20^ covariance models using their COVID-19 release 14.2 (http://rfam.xfam.org/covid-19) to highlight conserved non-coding RNA structures which are responsible for various stages of the viral life cycle. These include the frame shifting stimulation element and the pseudoknot necessary for the genome replication of SARS-CoV-2^21^. We also provide cross references to proteins from RefSeq, UniProt^22^ and the International Nucleotide Sequence Database Collaboration (INSDC); functional annotation from the Gene Ontology Consortium; and annotation of protein domains using InterProScan. These additional annotations are accessible via our region views and the gene and transcript tabs. We also created a genome browser track projecting the protein-domain annotations onto the genome to facilitate a genome-oriented view of the gene products including the non-structural cleavage products of ORF1a/ORF1ab.

The browser also displays community annotation of sites and regions using results co-ordinated by the UCSC genome browser. Additions to this annotation resource are open to all and done via a publicly available spreadsheet hosted by UCSC (http://bit.ly/cov2annots), the data from which is integrated periodically into the Ensembl browser. This is achieved via specialised code that uses Git workflows to convert the annotations into BigBed files that can be visualised on a variety of genome browsers (available freely at https://github.com/Ensembl/sarscov2-annotation).

We have integrated Oxford Nanopore sequencing primers (version 3) made available by the ARTIC network (https://artic.network/ncov-2019) to assist in sequencing the virus. Though mainly focused on the Oxford Nanopore MinION sequencer, some aspects of the protocol may be generalised to other sequencing platforms. The complete list of primers included is available on GitHub (https://github.com/artic-network/artic-ncov2019/blob/master/primer_schemes/nCoV-2019/V3/nCoV-2019.tsv).

Finally, we provide tracks to visualise problematic and caution sites, which result from common systematic errors associated with laboratory protocols and have been observed in submitted sequences^16^. Inclusion of these can adversely influence phylogenetic and evolutionary inference. Visualising these in the browser alongside the locations of primers and other community derived annotations helps determine how best to proceed with analyses of each these sites.

### Integration and engagement

The Ensembl COVID-19 resource features a newly designed landing page, which prioritises key views and data to help direct researchers into relevant sections of the site. To support expeditious data release, we have not made potentially time-consuming virus-specific modifications to our existing web codebase—such as showing a single nucleic acid strand and removing all mentions of exons— because we felt the data could be effectively understood without these changes. However, we have altered the vocabulary wherever possible and are reviewing feedback as we receive it.

Our COVID-19 resource is also integrated into the European COVID-19 Data Portal hosted by EMBL-EBI (https://www.covid19dataportal.org/). The portal enables searches across the multiple research outputs on COVID-19 including viral and human sequences; relevant biochemical pathways, interactions, complexes, targets and compounds; protein and expression data; and literature.

We have engaged our existing and new user communities using our blog and social media accounts to announce the release and updates to the Ensembl COVID-19 resource. We also highlighted the changes made to our gene annotation method to ensure the complete set of ORFs because these have been overlooked by other annotation tools.

## DISCUSSION

The swift spread of COVID-19 has highlighted the necessity for data resources to be prepared for rapid adaptation to developing outbreaks. Our development and release of the Ensembl COVID-19 resource leveraged our experience integrating thousands of genomes into the Ensembl infrastructure and supporting hundreds of thousands of users. The Ensembl COVID-19 browser provides a unique view on SARS-CoV-2 using our gene annotation method and variation data processed to focus on the highest confidence variants. Additionally, the Ensembl VEP and haplotype views enable the consequences of the variants to be assessed within the context of specific strains and geographical locations. The data is made accessible via the widely used Ensembl platform making it immediately familiar to a large userbase who may be able to repurpose existing software and browser knowledge to support their work during the pandemic and beyond.

When the COVID-19 pandemic hit, we had been working for several months to develop Ensembl Rapid Release (https://rapid.ensembl.org) to distribute annotated genomes within days of their annotation being completed. This experience proved useful in bringing the COVID-19 site to public release quickly. We have also demonstrated the flexibility of the Ensembl infrastructure and its value as a platform for research and discovery. Indeed, all of our pipelines and schemas worked seamlessly even though Ensembl was not designed to support RNA genomes and had not previously been used for viruses. The adapted gene annotation method, for instance, produced consistent annotation with ribosomal slippage correctly modelled and can be reused in the future. Similarly, the gene tree and alignment pipelines have been applied to the viral data with only minimal changes to parameters. We will continue to regularly update the site as new data emerges to support research into understanding the genomic evolution of this virus, identifying hotspots of genomic variation and enabling the rational design of future therapeutics, vaccines and policies well beyond the end of the current pandemic.

## ACKNOWLEDGEMENTS

We would like to thank the following people at the EMBL-EBI for their contributions to our resource and thoughtful discussions: Nick Goldman, Zamin Iqbal, Guy Cochrane, Rodrigo Lopez Sanchez, Conor Walker, Nadim Rahman, Jeena Rajan, Alexy Sokolov, Peter Harrison, Youngmi Park, Nicola Buso, Suran Jayathilaka, Anton Petrov, James Allen, Luca Da Rin Fioretto, Thomas Maurel and Vinay Kaikala.

This work was supported by the Wellcome Trust [WT108749/Z/15/Z] and the European Molecular Biology Laboratory. For the purpose of open access, the authors have applied a CC BY public copyright licence to any Author Accepted Manuscript version arising from this submission.

## AUTHOR CONTRIBUTIONS

N.H.D.S., R.D.F., P.F., A.F., K.L.H., S.E.H., F.J.M., M.R., A.D.Y. and D.R.Z. conceptualised the resource. N.H.D.S., S.E.H., F.J.M., M.R. and A.D.Y. contributed to the methodology. M.C., B.C., C.C., T.G., S.E.H., F.J.M., D.N.O., A Parker, A Parton, M.R., M.P.S., D.S., J.T. and A.T. developed the software. M.C., C.C., N.H.D.S., A.G., T.G., M.R., D.T. and A.D.Y. validated the data while C.C., T.G., K.L.H., S.E.H., M.R. and D.T. conducted formal analysis on the computed results. C.C., T.G., D.N.O. and D.T. helped with investigations of software and results. N.H.D.S. wrote the original draft of this manuscript and R.D.F., P.F., A.F., A.G., K.L.H., S.E.H., B.M., A Parker, M.R., D.T., S.J.T., A.D.Y. and D.R.Z. reviewed and edited it. A.W. created the visualisation for the resource landing page. N.H.D.S., R.D.F., P.F., K.L.H., S.E.H., M.R., S.J.T. and A.D.Y. supervised various aspects of the project. M.R. and A.D.Y. were involved in project administration and K.L.H., P.F., A.D.Y. and D.R.Z. acquired funds to support the project.

## DATA AVAILABILITY

The COVID-19 resource from Ensembl is available without restrictions at https://covid-19.ensembl.org. The reference genome assembly for SARS-CoV-2 with the accession GCA_009858895.3 was obtained from the European Nucleotide Archive (https://www.ebi.ac.uk/ena/browser/view/GCA_009858895.3).

## CODE AVAILABILITY

The selection of our code to convert CSV files into BigBed files is at https://github.com/Ensembl/sarscov2-annotation. The code relevant to processing SARS-CoV-2 variants in Ensembl is at https://github.com/Ensembl/ensembl-variation, the gene annotation pipeline is available at https://github.com/Ensembl/ensembl-annotation and the code used for comparative analysis is at https://github.com/Ensembl/ensembl-compara

